# Robust Characterization of Non-Invasive Post-Traumatic Osteoarthritis Mouse Model

**DOI:** 10.1101/2021.05.28.446049

**Authors:** Fazal-Ur-Rehman Bhatti, Yong-Hoon Jeong, Do-Gyoon Kim, David D. Brand, Karen A. Hasty, Hongsik Cho

## Abstract

**Objective:** Biochemical and molecular changes involved in the pathophysiology of post-traumatic arthritis (PTOA) have not been fully understood. This study used non-invasive mouse models to study biochemical, biomechanical and pain-related behavior changes induced in mice following repetitive mechanical knee loading. Mouse models were used to reflect the effects of the early stages of PTOA in humans.

**Methods:** Forty-eight twelve week old male mice were obtained for three groups: normal control without mechanical loading, trauma (24 hours after loading), and PTOA (early OA) groups. For the non-invasive PTOA mouse model, cyclic comprehensive loading (9 N) was applied on the left knee joint of each mouse. Biochemical and molecular changes induced by mechanical loading were analyzed after loading was completed. Blood and cartilage were collected and further examined using gene expression analysis. Grading of the tissue sections was completed using the osteoarthritis research society international (OARSI) scale. Biomechanical features of mechanically loaded knee joints were determined after 24 hours (Trauma) and three weeks (PTOA) post-mechanical loading sessions to examine the development of PTOA, respectively.

**Results:** The loaded left knee joint showed a greater ROS/RNS signal than the right knee that was not loaded. There was an increase in cartilage damage and MMP activity in the affected knee as the intensity of MabCII680 and MMP750 signal increased in the mechanical loaded joints as compared to unloaded control knee joints. There was also an increase in the difference of viscoelastic energy dissipation ability (tan δ) in PTOA. The OA score increased significantly in mechanically loaded knee joints.

**Conclusion:** This study showed that biomechanical, biochemical, and behavioral characteristics of the murine PTOA groups are significantly different from the control group. These results validate that the current mouse model can be used for translational studies to examine PTOA.

## Introduction

Injury to joint tissue leads to progressive damage to the articular joint leading to a condition known as post-traumatic arthritis (PTA) [1]. It has been reported that 20-50% of individuals subjected to joint trauma develop post-traumatic osteoarthritis (PTOA) that results in damage to not only cartilage tissue but also to other tissues that constitute the articular joint. Moreover, PTOA account for 12% of all osteoarthritis (OA) cases [2, 3].

An acute knee injury such as injurious trauma or excessive cumulative stress initiates biomechanical changes as well as biochemical cascades such as inflammation and metabolic imbalances of tissue turnover that can lead to secondary OA [4, 5]. This PTOA is characterized by multi-tissue damage such as articular cartilage, synovium, subchondral bone degradation. Evidence from both pre-clinical and clinical studies reveal that early intervention can alleviate these biochemical changes or altered joint biomechanics may slow the progression of joint degeneration [6].

Animal models are critical tools for PTOA research because they dramatically shorten the time required to develop PTOA. A previous study reported that a mature adult 3-6 month old C57BL/6J mouse is equivalent to a 20-30 year old human, a 10-14 month old middle-aged mouse is equivalent to a 38-47 year old human and an 18-24 month old mouse is equivalent to a 56-69 old human [7]. Mouse models are useful because they allow for investigation of specific genetic factors. In order to study mechanisms of PTOA initiation and progression that are relevant for human disease, it is important to utilize animal models that reproduce specific traumatic injury conditions seen in patients.

Animal models of PTOA can be broadly categorized in to invasive or non-invasive models. Invasive models involve surgical or chemical induction to develop OA. The invasive models require great deal of intervention and mostly represent acute phase of traumatic injury. Therefore, non-invasive mouse models have used in the past to mimic PTOA in humans to follow the progression of disease after traumatic injury [8, 9]. These models are developed by subjected a joint to repetitive mechanical load. The advantages of this model over invasive models are that these are non-invasive, no skin disruption and mimic early adaptive response to injury in humans. Mechanical loading of the mouse knee has been used as a model of PTOA and it can faithfully mimic injury conditions similar to that seen in early stages of human OA [9].

The pathology of PTOA can be characterized in three phases: the immediate phase following mechanical injury; the acute phase displaying apoptosis and inflammation; and the chronic phase characterized by dysfunction and pain of joint [10]. The biochemical changes occurring after trauma depends mainly whether the impact is either of low or high amplitude [11]. The most common irreversible biochemical change that occurs is the breakdown of cartilage matrix due to disruption of proteoglycans that expose the underlying collagens that maintain the stability of cartilage matrix. In particular collagen type-II (COL2A1) is the main collagen affected in hyaline cartilage [12]. This, in turn, causes necrosis of chondrocytes residing within the cartilage matrix. At the molecular level various factors responsible for the catabolism of cartilage tissue include production of matrix metalloproteinases (MMPs), reactive oxygen species (ROS) and inflammatory cytokines [13–15]. ROS are released from storage in the mitochondria after excessive mechanical stress in the knee joint. Release of these destructive metabolites can result in chondrocyte death and matrix degradation [5].

Although conventional mechanical loading methods have been utilized to study the pathophysiology of PTOA, past studies have failed to include a comprehensive biochemical and molecular characterization. In a previous study, we developed an *in vivo* detection technique for visualizing early damage to cartilage using a fluorescence labeled monoclonal antibody (Mab) that binds to type II collagen (CII) [16]. This method has the advantage of detecting damaged cartilage in the mouse knee joint.

A functional characterization such as pain-related behavior, along with previous conventional methods of studying PTOA is essential for translation of preclinical studies to clinical OA therapy. Therefore, we developed this PTOA mouse model to characterize all facets of arthritis from the macroscopic to the molecular level. Specifically, we provide behavioral data related to the OA pain mechanism that has not been addressed in previous studies. In brief, we discuss the behavioral, biochemical and molecular changes that occur following non-invasive repetitive mechanical loading of the mouse knee joint.

## Materials and Methods

### Animals

A total of forty-eight 12 week old (n=16 per group) CBA-1 male mice were used in this study. Mice were randomly divided into three test groups (Table 1): normal control without mechanical loading, Trauma (the immediate phase following mechanical injury which involved the cell apoptosis and joint inflammation), and PTOA (early OA) group. Mice were provided with a pathogen free environment with adequate food and water *ad libitum.* All animal protocols and experimental procedures were approved by the Institutional Animal Care and Use Committee (IACUC) at the University of Tennessee Health Science Center.

**Table 1.**
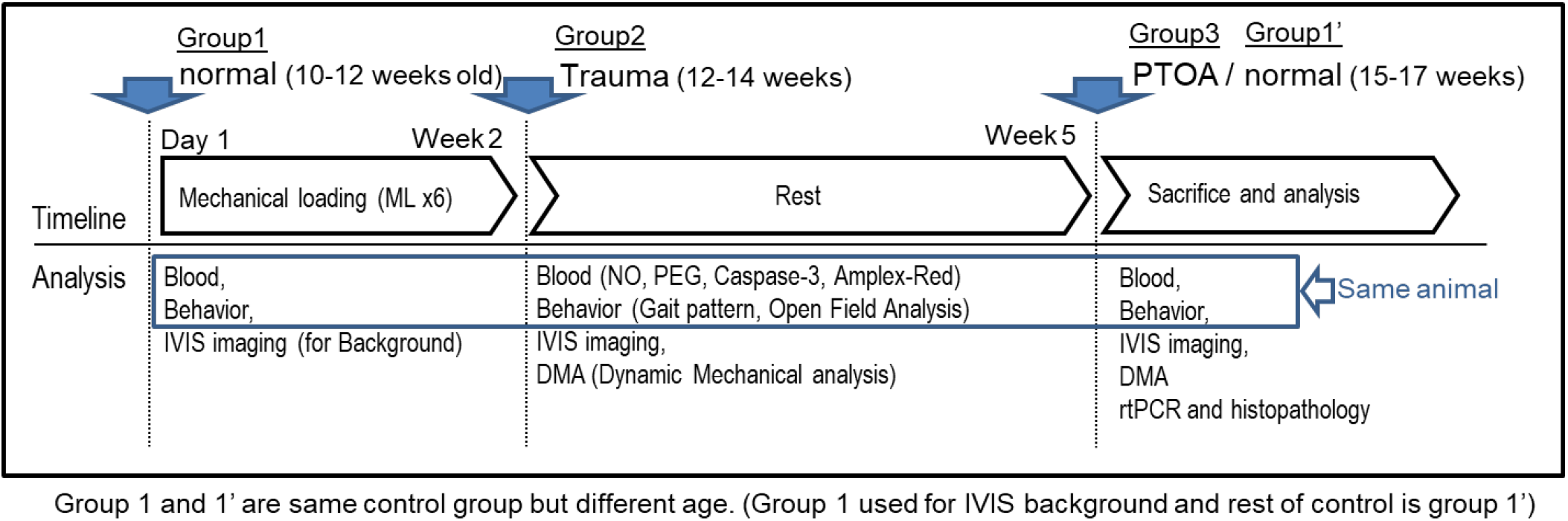
Schematic representation of the study. Groups represent the control (Group 1) and experimental groups (Group 2 and 3). Group 1 is the control group that was not subjected to mechanical load. Group 2 is classified as trauma group that shows acute tissue damage 3-week post mechanical load period. Group 3 represents PTOA mouse model showing chronic tissue damage 10-12 week post mechanical load period.

### Non-invasive post-traumatic osteoarthritis (PTOA) mouse model

The PTOA mouse model was developed as described previously with slight modifications [17–19]. Briefly, mice were continuously anesthetized throughout the procedure using 2% isoflurane. The left leg of each mouse was positioned within the ElectroForce® 3200 (Bose Corp., Minnesota, USA) using a custom-made mechanical loading jig. The knee joint was positioned with the proximal tibia resting in the upper cup and the dorsiflexed ankle inserted into the bottom cup. Each loaded knee joint received 40 cycles of compressive loading at 9 N, every other day for nine successive sessions. The left leg of control animals also underwent the same procedure for the same time interval except that no compressive loading was applied. Animals were left with adequate food and water *ad libitum* following each mechanical loading session and monitored for behavior. The mechanical loading apparatus was calibrated before each loading session. Loading data was collected using WinTest software (Bose Corp., Minnesota, USA).

### Dynamic mechanical analysis (DMA) of the knee joints

The femur and tibia were separated, and soft tissues were completely removed. Tibia was glued on the jig of ElectroForce® 3200 instrument (supplement Fig 3). The specimen was secured by preloading at 1 N on the loading jig. Next, compressive cyclic displacement (0.01 mm ± 0.0025 mm) at the range of 0.5 to 3 Hz was applied as described previously [20, 21]. The corresponding force ranged between 0 and 0.5 N. Displacement was controlled by a transducer with 15 nm resolution. The cyclic force and displacement were measured to obtain dynamic elastic (storage) (K’) and viscous (loss) (K”) stiffness. Dynamic complex stiffness (K*) was computed using the following equation: K*= K’+iK”. The K’ and K” of bone represent its abilities to store and dissipate the dynamic loading energy, respectively. Viscoelastic tangent delta (tan δ), which represents loading energy dissipation, was computed by K”/K’. As such, the tan δ accounts for the relative capacity of dynamic energy dissipation [53,54].

### Gait analysis

Gait analysis was performed as described previously with slight modifications [22]. Briefly, hind and fore paws were stained with black and red nontoxic ink, respectively. Each animal was allowed to walk along a 100 x 10 x 15 (L x W x H cm) runway lined with white paper ending in a dark enclosed escape box. Animals were trained and acclimatized prior to recording the gait pattern.

### Open-field analysis

Open field analysis was performed using a custom-grade Open Field Maze instrument (30 x 20 x 20 cm^3^) [23]. The animals were acclimatized to the Open Field Maze before the actual readings were made. Briefly, each animal was placed in the center of instrument and was allowed to move freely. The movement of each animal was recorded for 10 min. Any type of disturbance was avoided during the experiment. A five-minute segment was analyzed manually using CowLog 3.0.2 software to determine the total travel distance and rearing activity for each animal [24].

### Optical image scanning

Optical image scanning was performed as described previously [16]. At the conclusion of mechanical loading sessions, each mouse was injected retroorbitally with 80 μl of solution containing a 1:1 mixture of MMPSense 750 FAST Fluorescent Imaging Agent (MMP750) (PerkinElmer, Waltham, MA) and monoclonal anti-type II collagen antibody that had been coupled with 680 dye (MabCII680) using XenoLight CF680 Fluorescent Rapid Antibody Labeling Kit (Caliper Life Sciences, Waltham, MA).

The monoclonal anti-type II collagen antibody was provided by the VA Program Project Scientific Core at the VA Medical Center (Memphis, TN). The generation and characterization of the anti-collagen type II monoclonal antibodies (MabCII) are described previously [52]. Briefly, an immunoassay was performed to measure binding and specificity of these antibodies for CII purified from mouse cartilage. The results showed that monoclonal anti-CII antibody (MabCII) (E4), the antibody chosen for this study, had the strongest immune reactivity.

24 hours after injection with the probe cocktail, mice were imaged under anesthesia after using an IVIS In vivo Imaging System (IVIS® Lumina XR System, Perkin Elmer, Hopkinton, MA). The excitation/ emission wavelengths for MMP750 and MabCII680 were 745 nm/ 800 nm and 675 nm/ 720 nm, respectively. Fluorescence was quantified using Living Image 4.0 software to calculate the flux radiating omni-directionally from the region of interest (ROI) and graphed as radiant efficiency (photons/s/cm^2^/str)/ (μW/cm^2^). The standardized ROI for knee fluorescence was obtained by capturing the same area for each mouse. The background signal from other tissues, such as muscle and skin tissue, was measured and subtracted from each articular reading.

The level of reactive oxygen species (ROS) and reactive nitrogen species (RNS) in the knee joint was measured after the first mechanical loading session. This time point was chosen because ROS are measureable immediately after loading and because after the multiple sessions the ROS/RNS signal can be detected systemically. Both knees were injected with 25 mg/Kg body weight L-012 sodium salt (TOCRIS, Bristol, UK) through intra-articular route, and scanned immediately after 1 min by IVIS to determine the chemiluminescent signal [25]. The ROI was selected and calculated as described earlier.

### Collection of blood and isolation of cartilage tissue

After the optical image scanning, blood was collected from each animal through retro-orbital method using a sterilized heparinized capillary tube (Fisher Scientific, Hampton, NH). Serum was collected by centrifugation of blood at 3,000 rpm for 5 min. Serum was stored at −70°C until further use.

Following blood collection, the animal was euthanized and the entire knee joint was isolated and dissected. The knee joint was immediately transferred to RN*Alater™* Stabilization Solution (ThermoFischer Scientific, Waltham, MA). Under sterile conditions, the soft tissue was removed from the bones and the cartilage tissue was isolated from femoral heads and tibial plateau. The isolated cartilage tissue was processed immediately for the extraction of RNA as explained later.

### Total nitrate/nitrite (NO) Assay

Serum was analyzed for nitrates and nitrites using the Nitrate/Nitrite Fluorometric Assay Kit according to the manufacturer’s instructions (Cayman Chemical Company, Ann Arbor, MI). Fluorescence signal was read at an excitation wavelength of 375 nm and emission wavelength of 417 nm using SpectraMax M5 microplate reader (Molecular Devices, Sunnyvale CA). The total NO was interpolated from the nitrate standard curve.

### Caspase3 assay

Caspase3 activity in serum was analyzed using the EnzChek® Caspase-3 Assay Kit (Molecular Probes™, Eugene, OR) according to the manufacturer’s instructions. Fluorescence was read at excitation and emission wavelengths of 496 and 520 nm, respectively, using SpectraMax M5 microplate reader. The amount of fluorescence was used to determine the level of Caspase3 activity in serum.

### Amplex Red assay for H_2_O_2_

The level of H_2_O_2_ was measured in serum using the Amplex® Red Hydrogen Peroxide/Peroxidase Assay Kit (Thermo Fisher Scientific, Waltham, MA). Fluorescence was read at excitation and emission wavelengths of 571 and 585 nm, respectively, using SpectraMax M5 microplate reader.

### Prostaglandin E2 (PGE_2_) assay

Serum was analyzed for PGE_2_ concentration with an Enzyme Immunoassay kit (Cayman Chemical Co., Inc., Ann Arbor, MI). The microtiter plate was read at a wavelength of 405 nm using a SpectraMax M5 microplate reader.

### Gene expression analysis

RNA was extracted from the cartilage tissue using the GeneJET RNA Purification Kit (Thermo Fisher Scientific, Waltham, MA). cDNA was prepared using 0.5 μg RNA by TaqMan® Reverse Transcription Reagents (Thermo Fisher Scientific, Waltham, MA). Semi-quantitative gene expression (qPCR) was performed using a TaqMan™ Gene Expression Assay (Thermo Fisher Scientific, Waltham, MA) for the following genes: Aggrecan (ACAN), Collagen type II alpha (COL2A1), Endothelial PAS domain-containing protein 1 (EPAS1), Matrix metallopeptidase 13 (MMP13), Smad Nuclear Interacting Protein 1 (SNIP1), Interleukin 1 Beta (IL1β) and Tumor necrosis factor alpha (TNFα). Beta-Actin (β-Actin) served as an internal control. qPCR was performed on a LightCycler®480 (Roche, Basel, Switzerland). Data was analyzed using LightCycler® 480 software (Roche, Basel, Switzerland).

### Histology

The entire knee joint was fixed in 10% formalin solution, decalcified with Richard-Allan Scientific™ Decalcifying Solution (ThermoFischer Scientific, Waltham, MA) and embedded in paraffin. A total of twenty-five histological sections were analyzed for across the entire joint to evaluate cartilage damage and synovitis. The sections were stained with Hematoxylin and eosin stain (H&E) or Safranin-O/ Fast Green stain.

### Modified OA and synovitis scoring system

Tissue sections were graded using the osteoarthritis research society international (OARSI) scale with slight modifications [26]. The advantage of the OARSI system is that it allows the tissue sections to be scored based on grade and stage of disease progression. Thus, a higher score represents a high level of biological deterioration and disease extent. Therefore, it is reliable method to differentiate between early-to latest-age OA. We further modified the procedure to assess the whole knee joint from an individual animal and to sum up the average histological score from each representative group.

Briefly, the whole knee joint was divided in to four groups as follows: medial femoral condyle, medial tibial plateau, lateral femoral condyle and lateral tibial plateau. Each area was graded based on hypocellularity, Safranin-O staining, surface regularity and structure. The mean from each area was calculated and summed up with other areas to yield sum of average histological score for each group of animals. Synovitis was graded as described previously with slight modifications [27]. Briefly, synovitis score was based on enlargement of synovial lining, density of cells and inflammation.

### Statistical analysis

Statistical significance was determined by the Student’s t tests. A *P*-value ≤ 0.05 was considered statistically significant. GraphPad Prism v.5.00 for Windows (GraphPad Software, USA, http://www.graphpad.com) was used to perform statistical analysis.

## Results

### Mechanical loading induced oxidative stress on knee joints

Initially we studied the induction of oxidative stress following mechanical loading of the knee joint. Oxidative stress was obvious following the first mechanical loading session (Figure 1). The Bioluminescence signal increased significantly in the unloaded control right knee joint as compared to mechanical loaded left knee joint (4.46 ± 0.68 ×10^3^ ROI of ROS in control knee vs 338.25 ± 88.33 ROI of ROS in ML knee, *P* < 0.0001). This indicated the mechanical load increases both ROS and RNS. We also confirmed the binding of MabCII680 to the damaged cartilage tissue in explant pig cartilage tissue and mouse knee joints (Figure 2). MabCII680 specifically binds to the damaged cartilage owing to the exposed COL2A1. Furthermore, examination of mouse knee joints demonstrated that the intensity of MabCII680 and MMP750 signal increased in the traumatic and mechanically loaded joints relative to unloaded control knee joints (1.47 ± 0.43 ×10^7^ ROI of MabCII680 in normal vs 19.34 ± 2.49 in Trauma vs 23.52 ± 4.85 in PTOA knee joint, *P* < 0.001). The MMP750 signal was significantly increased in the traumatic joint relative to unloaded control knee joints but the signal decreased over time in the posttraumatic period (6.38 ± 1.19 ×10^5^ ROI of MMP750 in normal vs 37.94 ± 3.42 in Trauma vs 13.78 ± 5.51 in PTOA knee joint, *P* < 0.001).

**Figure 1.**
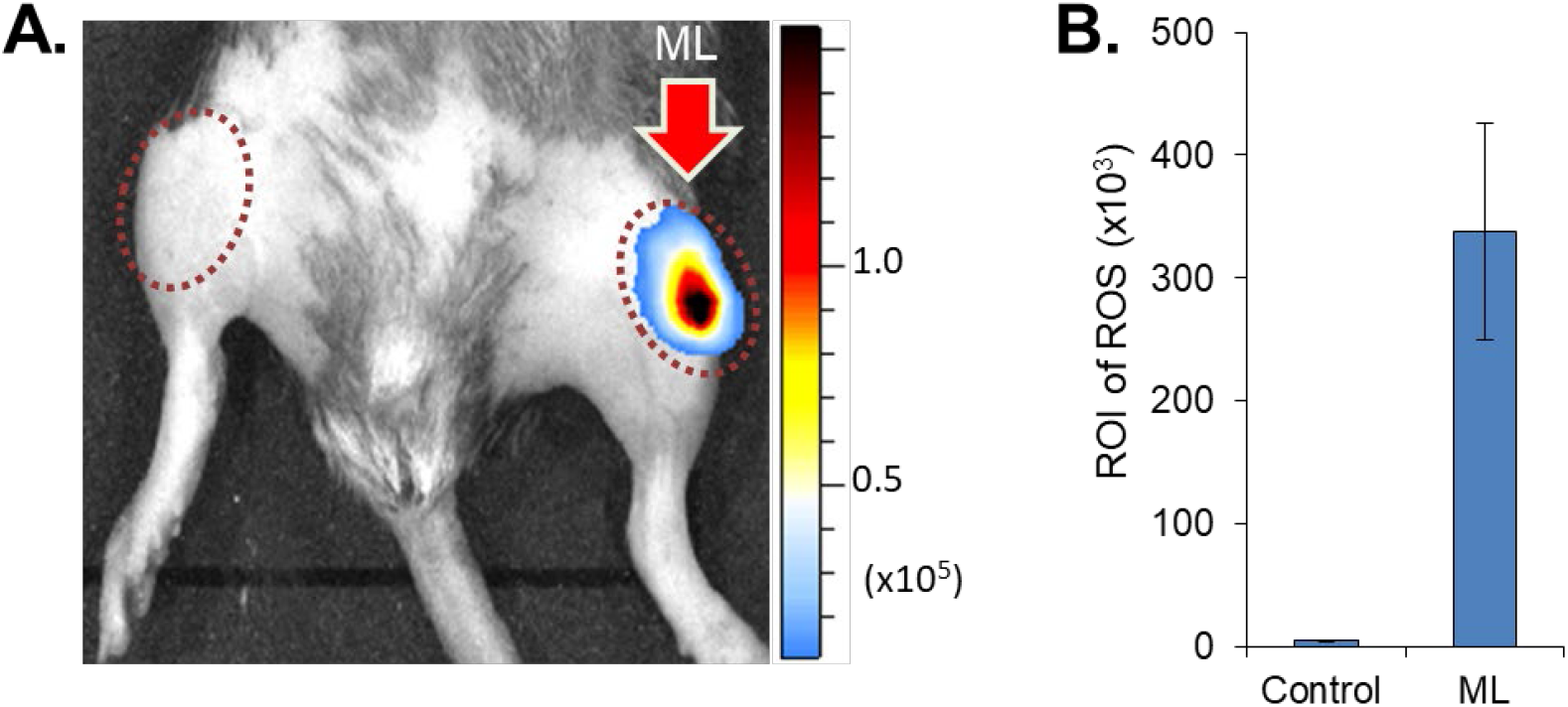
Comparison between mechanical loaded (ML) and unloaded knee joint. (A) IVIS image showing the acute damage to knee joint following 1st mechanical load session. Observe high level of ROS-bioluminescence in the ML knee. (B) Graphical representation of ROS-bioluminescence in the region of interest (ROI). Data is represented as mean ± SD. (*** *p* < 0.001).

**Figure 2.**
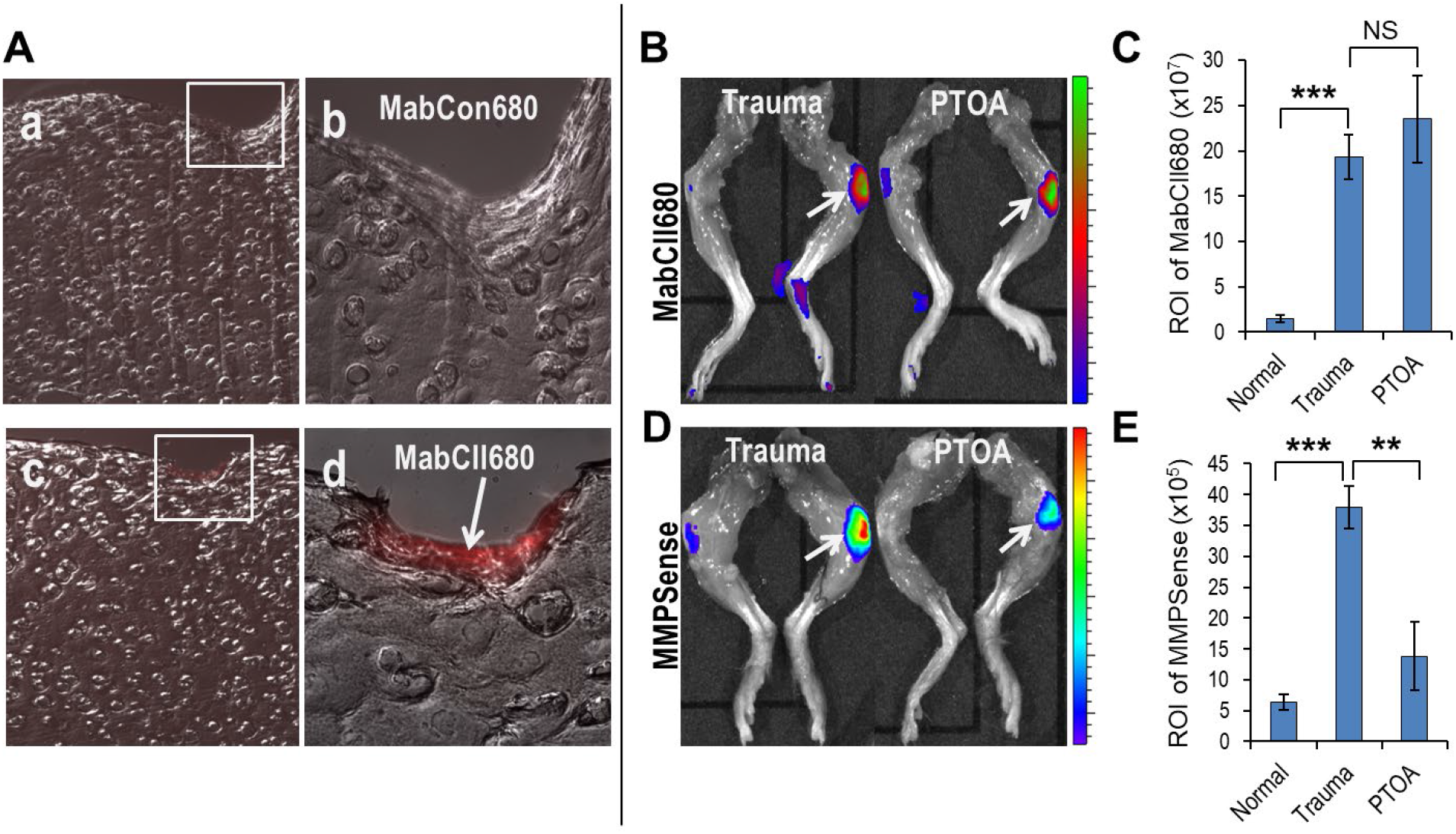
Assessment of cartilage damage and MMPs level in trauma and PTOA groups. (A) Histological sections showing specificity of MabCII680 (lower panel) to the exposed type-II collagen in the damaged tissue as compared to control MabCon680 at low and high magnification (B, C) IVIS image and graphical representation of cartilage damage in trauma and PTOA knee joints. (D, E) IVIS image and graphical representation of MMPs level in trauma and PTOA knee joints. Data is represented as mean ± SD. **p < 0.01 and ***p < 0.001.

### Effect of mechanical loading on biomechanical properties

Alterations in the biomechanical properties are one of the early signs of PTOA [28]. Therefore, we determined the biomechanical features of mechanically loaded knee joints under conditions of acute trauma (Trauma group) and 3 weeks after mechanical loading to examine the development of PTOA (PTOA group). Increases in the difference of viscoelastic tan δ (1.0 ± 0.128 in normal vs 3.326 ± 0.331 in Trauma vs 2.817 ± 0.311 in PTOA, *P* < 0.0001) was observed relative to control unloaded knee joints (Figure 3). In this result, we found a change in the mechanical properties of cartilage and subchondral bone based on the increased value in Viscoelastic tangent delta (tan δ).

**Figure 3.**
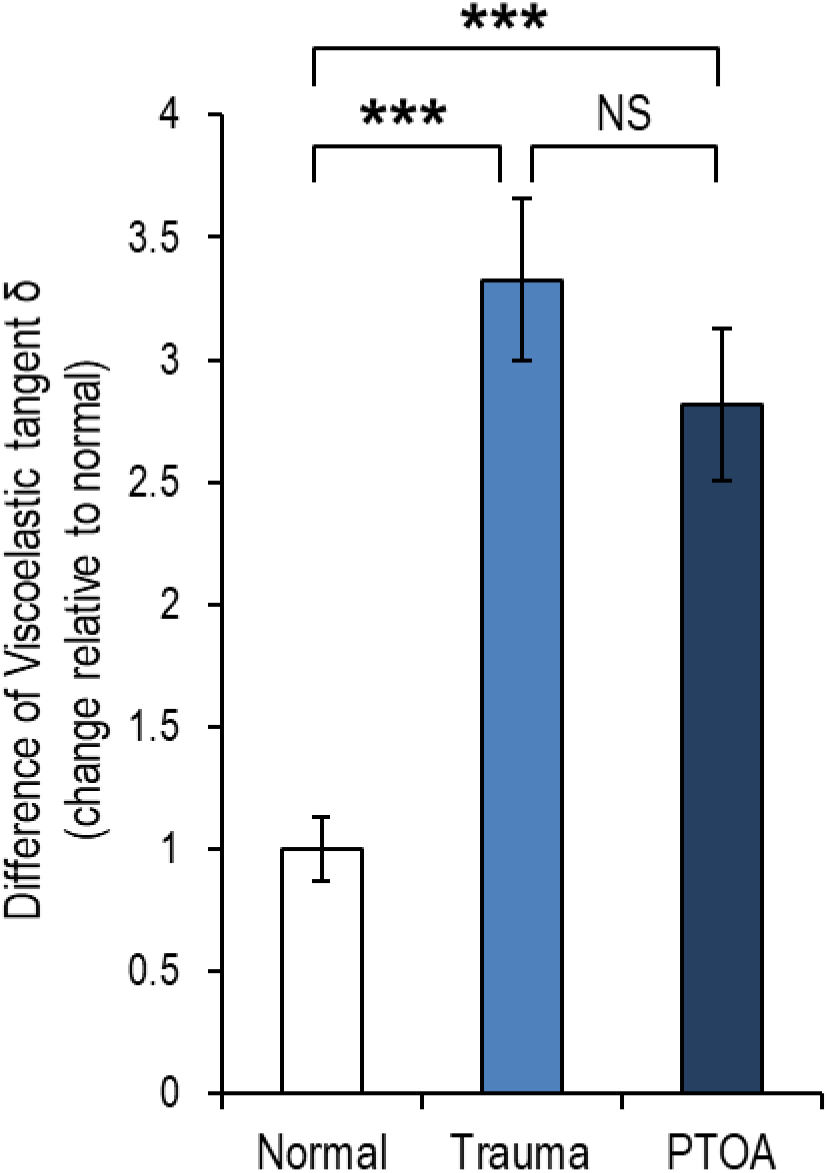
Dynamic Mechanical Analysis (DMA). Difference of viscoelastic tan delta (δ) on mechanical loaded knee (Trauma vs PTOA) vs contralateral knee. Data is represented as mean ± SD. ***p < 0.001.

**Figure 4.**
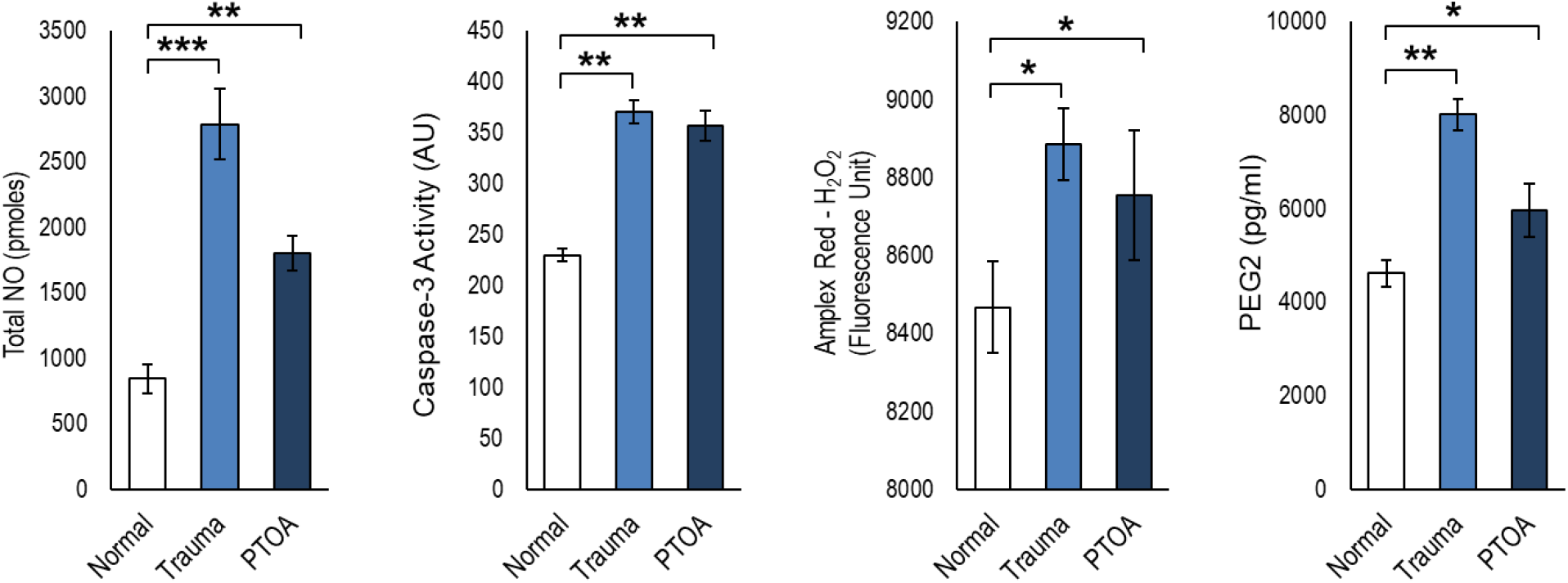
Serum biochemical analyses of trauma and PTOA to control unloaded normal knee joint. Level of (A) Total NO, (B) Caspase-3 activity, (C) Amplex Red H_2_O_2_ and (D) PEG2 in sera. Data is represented as mean ± SD. *p < 0.05, **p < 0.01 and ***p < 0.001.

### Biochemical changes accompanied by mechanical loading

We took a comprehensive approach to our mouse model of PTOA. We not only analyzed the gross pathological changes in joint architecture and the resultant change in function, but also designed our study to dig deeper into the biochemical events that may be at least, in part, responsible for driving the disease process. We first measured total Nitric Oxide and found a significant increase in total NO in our study in mice subjected to mechanical loaded as compared to the untreated group (2790.77 ± 273.12 pmoles vs 845.67 ± 110.94 pmoles, *P* = 0.0003) (Figure 3A). [29, 30].

In addition to NO, we measured Caspase-3 levels in the joint to determine if apoptosis might be driving pathology. [31, 32] We observed that mechanical loading caused a significant increase in caspase-3 activity as compared to untreated control (370.65 ± 11.09 fluorescence unit vs 230.04 ± 6.37 fluorescence unit, *P* < 0.0001) (Figure 3B).

Another mediator of oxidative stress that can result in destruction of tissues is hydrogen peroxide (H_2_O_2_). [33]. We observed that H_2_O_2_ levels were significantly higher in the mechanically loaded group relative to the untreated control group (8884.88 ± 92.06 fluorescence unit vs 8468.24 ± 118.59 fluorescence unit, *P* < 0.0001) (Figure 3C).

We also opted to measure PGE2 levels that are reported to exert catabolic effects under mechanical stress [34, 35]. We observed that mechanical loading caused significantly high PGE2 level in mechanically loaded mice as compared to untreated control (8015.62 ± 36.72 pg/ml vs 4627.50 ± 86.66 pg/ml, *P* < 0.0001) (Figure 3D).

### Mechanically induced gene expression patterns

The imbalance between anabolism and catabolism leads to pathogenesis of OA [36]. We studied the expression of both anabolic and catabolic genes. The expression of major proteoglycan (ACAN) and collagen (COL2A1), which are markers of anabolism were both reduced after mechanical loading relative to untreated control; ACAN (0.58 ± 0.03-fold vs. 1.00 ± 0.004-fold, *P* < 0.0001) and COL2A1 (0.40 ± 0.10-fold vs. 1.01 ± 0.28-fold, *P* = 0.0245) (Figure 5A and B). In addition, the expression level of SNIP1, an inhibitor of NFκB signaling decreased in mechanical loaded mice relative to untreated controls (0.24 ± 0.10-fold vs. 1.01 ± 0.27-fold, *P* = 0.0108) (Figure 5E).

**Figure 5.**
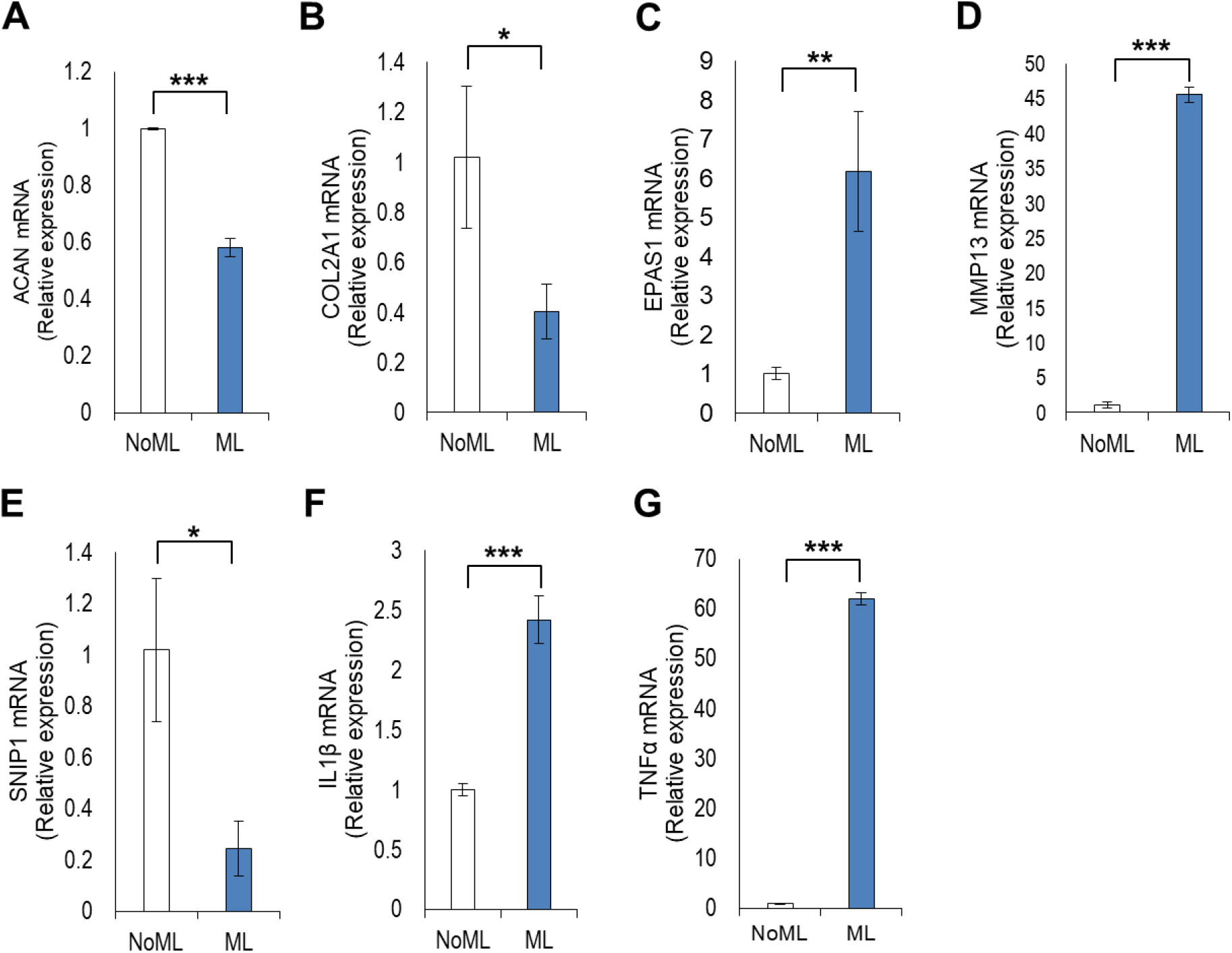
Gene expression analysis in cartilage tissue of mechanical loaded knee to control unloaded knee joint. Gene expression level of (A) ACAN, (B) COL2A1, (C) EPAS1, (D) MMP13, (E) SNIP1, (F) IL1β and (G) TNFα. Data is represented as mean ± SD. *p < 0.05, **p < 0.01 and ***p < 0.001.

Our analysis revealed that genes responsible for cartilage catabolism increased after mechanical loading of the knee joint. EPAS1, a gene that encodes hypoxia-inducible factor (HIF-2α) increased significantly in the mechanically loaded group relative to the untreated group (6.17 ± 1.52-fold vs. 1.00 ± 0.14-fold, *P* = 0.0043) (Figure 5C). In addition, the expression of MMP13, a gene encoding an enzyme that degrades the cartilage matrix also increased significantly in the mechanically loaded group relative to the untreated group (45.57 ± 1.11-fold vs. 1.04 ± 0.44-fold, *P* < 0.0001) (Figure 5D). The inflammatory cytokines IL1β and TNFα were both enhanced significantly after mechanical loading as expected relative to untreated controls; IL1β (2.41 ± 0.19-fold vs. 1.00 ± 0.05-fold, *P* = 0.0003) and TNFα (62.04 ± 1.21-fold vs. 1.00 ± 0.11-fold, *P* < 0.0001) (Figure 5F and G). These inflammatory cytokines not only result in knee joint inflammation but also drive MMP13-mediated degradation of cartilage tissue.

### Mechanical loading of the knee joints increased OA and synovitis score

Mechanical knee joint loading significantly impacted cartilage integrity (Figure 6A and B). The medial compartment was especially affected, possibly because the medial side bears more weight. The most prominent features in osteoarthritic joints were a pronounced reduction in Safranin-O staining and a slightly irregular joint surface (Figure 6C and D). We also observed synovitis in knee joints subjected to mechanical loading (Figure 6E and F). The synovial membrane of osteoarthritic knees was highly inflamed relative to untreated controls. Concomitantly, OA scores increased significantly in mechanically loaded knee joints. Taken together, the histopathological observations clearly demonstrated signs of PTOA as evident by cartilage damage and inflammation.

**Figure 6.**
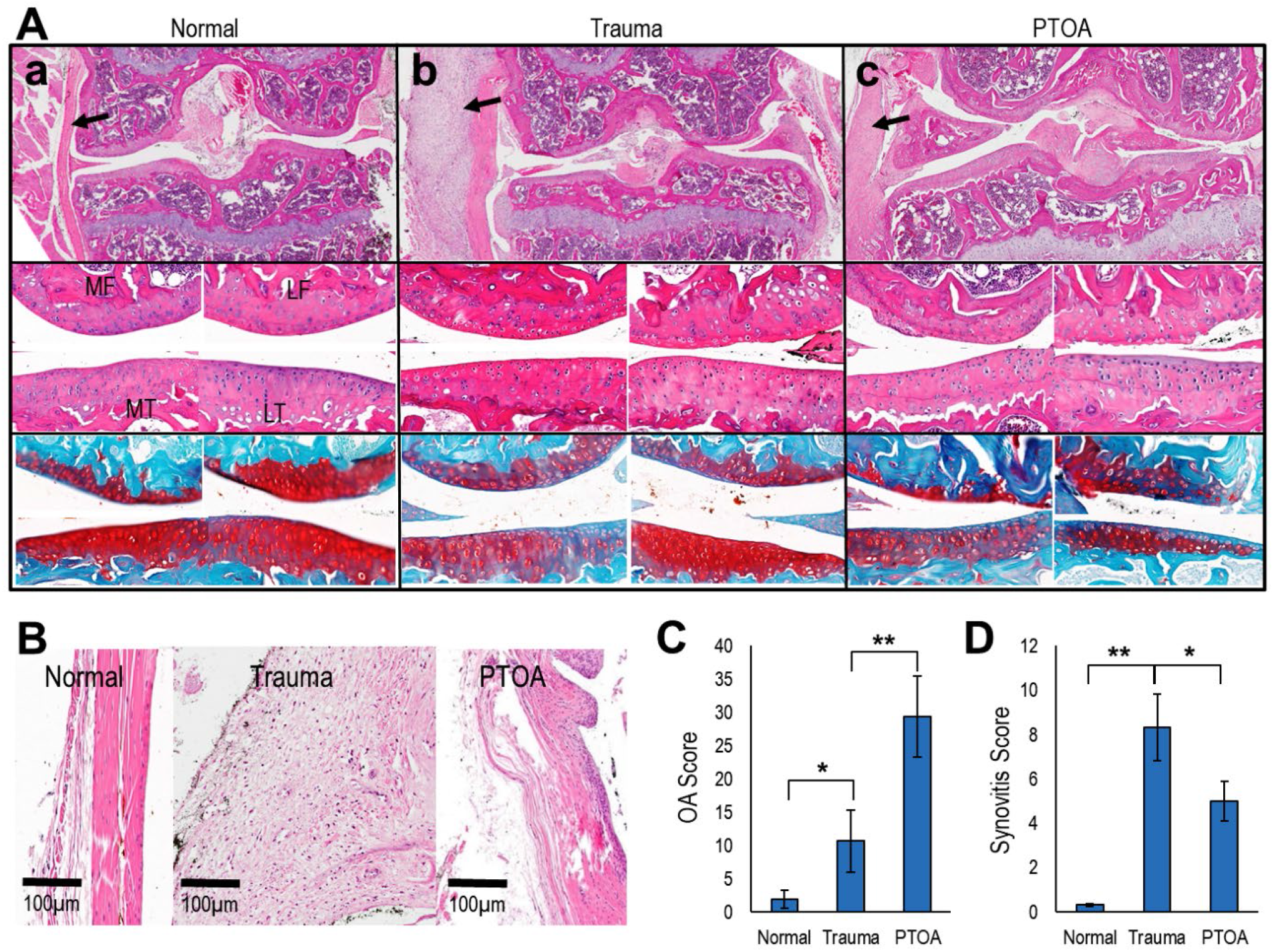
Histopathology of knee joints, synovial membrane and scoring according to OA and synovitis score. (A) Representative H&E stain of whole knee joint (upper panel). Observe inflamed synovial membrane as indicated my b;ack arrows. Magnified images of the same knee joint showing four distinct regions of the knee joint (middle panel). Cell distribution is affected by cartilage damage. Safranin-O/ Fast Green stain of the same knee joint (lower panel). Proteoglycan loss is obvious in trauma and PTOA groups. Symbols: MF = medial femur, LF = lateral femur, MT = medial tibia and LT = lateral tibia. (B) Representative H&E stain of synovial membrane. (C) OA score and (D) Synovitis score. Data is represented as mean ± SD. *p < 0.05 and **p < 0.01.

### Altered gait and locomotory behavior in mechanical loaded mice

In order to gain a perspective on the functional consequences of joint loading, we subjected mice to gait analysis. We observed that mechanically loaded animals displayed a highly abnormal synchronous gait pattern relative to control alternate limb movements in control mice (Figure 7A). Spatial variables including stride length, sway length (step width), and stance length describe the geometric position of the paw prints during locomotion. Rodents tend to have symmetric gaits, where left and right limb foot-strikes (for either the fore or hind limbs) are spaced at approximately 50 % of the gait cycle in time and equidistant in space (step length is about 50 % of stride length).

**Figure 7.**
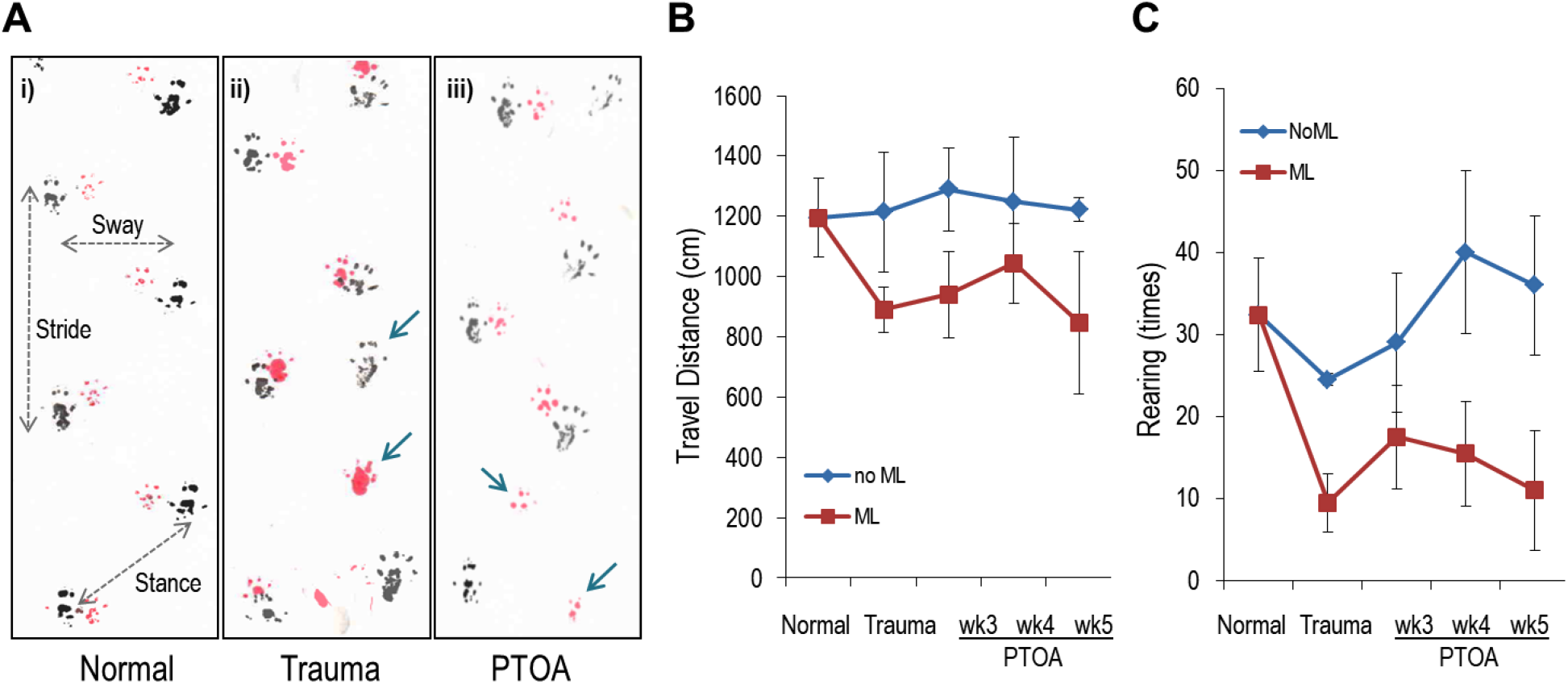
Behavior analyses of control and experimental animals. (A) Gait analysis. Observe the difference in stance, stride and sway among the groups. (Blue arrow indicate the missing step / asymmetric gait patterns). Open field analysis between unloaded control and mechanical loaded mice. (B) Travel distance and (C) Rearing activity. Data is represented as mean ± SD.

Mechanical loading (both Trauma and PTOA group) reduced the stride length (−9.5% in Trauma and −4.9% in PTOA group), sway length (−6.1% in Trauma and −1.4% in PTOA group), and stance length (−12.3% in Trauma and −1.6% in PTOA group). Missing step and asymmetric gait patterns occurred with the unilateral injury. The missing step rates were increased in both Trauma (9.36%) and PTOA (4.64%) groups when compared to normal control. (figure 7A and Table 2).

**Table 2.** Gait Analysis using mice foot print (Fig. 7. A). Missed-step rates were calculated by number of missed steps divide by total number of steps.

|  | Normal (N) | Trauma | PTOA | N v Trauma | N v PTOA |
| --- | --- | --- | --- | --- | --- |
| Stride (cm) | 7.066 ± 0.163 | 6.400 ± 0.167 | 6.717 ± 0.264 | p= 0.001 | p= 0.031 |
| Sway (cm) | 3.533 ± 0.103 | 3.317 ± 0.194 | 3.483 ± 0.223 | p= 0.035 | p= 0.245 |
| Stance (cm) | 4.183 ± 0.160 | 3.667 ± 0.175 | 4.117 ± 0.204 | p= 0.003 | p= 0.142 |
| Missing step (%) | 0.35% | 9.36% | 4.64% |  |  |

To investigate immobility in the PTOA mouse model, we performed open field analysis. Open field analysis revealed a significant reduction in the total distance travelled (reduced average 26.69% in Trauma group and 30.68% in PTOA group when compared to Normal control) and rearing (reduced average 61.23% in Trauma group and 67.44% in PTOA group when compared to Normal control) following mechanical loading (Figure 7B, C). Evaluation of locomotor activity in the PTOA mice showed that the mean moving distance and rearing counts were significantly different from normal mice. (Figure 7B, C).

## Discussion

Mechanical injury is used to develop a murine model of PTOA to study the etiology of early OA in humans [9]. A previous study investigated the lesions resultant from applied mechanical knee joint loading on eight-week old male mice [37]. As in our study, a force of 9N resulted in PTOA. However, the focus of their study was primarily directed towards the characteristization of joint lesions. We further characterized our murine PTOA model using a comprehensive approach including mechanical, biochemical, molecular and functional changes. One of the main differences between the two studies is that the previous study used the inbred mouse strains LGXSM-6 and LGXSM-33 which differ in their ability to regenerate knee cartilage and also differ in the development of PTOA following destabilization of medial meniscus (DMM) [38]. In the present study, we utilized CBA-1 mice that are known not to develop any evidence of spontaneous OA and therefore serve as an effective control [39].

A recent study on chondrocytes and mouse femoral head explants showed that excessive mechanical loading results in the production of ras-related C3 botulinum toxin substrate 1 activation (RAC1) that enhances ROS, resulting in the activation of the Nuclear Factor kappa-light-chain-enhancer of activated B cells (NF-κB) pathway, ultimately resulting in the production of catabolic enzymes leading to OA [40]. All these changes were attributed to the induction of gremlin-1 gene. We found that the production of ROS/RNS was evident as early as the first mechanical loading session. The intensity of the bioluminescence signal was higher after mechanical loading and declined over time (data not shown). At a biochemical level, we observed increases in the levels of total NO and H_2_O_2_ in mechanically loaded mice. Therefore, we propose that ROS/RNS production plays a significant role in early PTOA and suggest that studies focused on the role of oxidative stress in the development of PTOA in murine model should rely on early time points for ROS/RNS measurements.

We previously developed a monoclonal antibody (MabCII680) that binds only to exposed type II collagen in damaged cartilage [16]. We have also established that inflammation can be quantified through measurement of matrix metalloproteinase enzyme activity using the MMP750 probe [41]. In this study, we found increases in the signal intensity of both MabCII680 and MMP750 following mechanical loading. This finding suggests that cartilage damage and inflammation may be a consequence of mechanical load. In addition, we found that mechanical stress also compromises the biomechanical properties of articular cartilage. It is well known that structural and biochemical changes are associated with the pathogenesis of OA [42, 43]. The changes observed in the viscoelastic properties in this study can be explained due to an increase in the water content of articular cartilage matrix with a concomitant decrease in proteoglycan levels and in type II collagen content [44, 45]. Interestingly, a previous study demonstrated changes in viscoelastic properties prior to any histologic signs of OA [46]. In contrast, we observed changes in the biomechanical properties along with histological changes.

Inflammation and apoptosis are well-established key players in the generation of OA [47, 48]. We found mechanical loading significantly enhanced joint inflammation. The PTOA mouse model demonstrated an increased serum production of PGE2 and also an increase in the gene expression levels of IL1β and TNFα. Furthermore, Caspase-3 activity levels in serum increased following application of a mechanical load. These data indicate that serum markers and articular cartilage gene expression analyses can both participate in the development of OA.

Histopathological analysis remains the gold-standard in establishing the presence of OA in animal models [49]. Our histological analyses clearly showed signs of osteoarthritic changes occurring following mechanical loading. Particularly, we found that the medial compartment is primarily affected as a result of this area being the surface bearing the most weight. A significant loss of proteoglycans and enhanced synovitis score were the main features observed in this compartment. Meniscal damage was also prominent in osteoarthritic knee joints. Thus, structural changes were vividly established in our murine PTOA model, and these changes were consistent with the biomechanical, biochemical and molecular alterations observed earlier.

We also evaluated the outcomes of structural and biochemical changes in terms of the functional ability of mice affected by PTOA [50]. In our study, it was obvious that gait pattern and locomotor activity were severely affected in PTOA mice. These observations highlight the utility of functional outcome measurements in the PTOA mouse model. It is noteworthy that our observations correlated with a previous study showing that mechanical loading of knee joints with 9N force produces lesions in ipsilateral and contralateral joints [51]. The authors indicated that the progress and worsening of lesions corresponded to the nociceptive behavior in loaded animals. Moreover, it was emphasized that 9N force was more appropriate for longitudinal pharmacological studies due to mild joint damage. Our study not only is in conformity with these previous findings but also provides insight into the behavioral changes due to the application of the 9N force for mechanical loading. It is vital to apply an optimal force in order to study the desired stage of disease. Thus, our model clearly showed that the application of 9N according to the method explained in our study will allow us to measure functional changes associated with the application of a mechanical load.

Post-loaded mice clearly show lesions in the articular cartilage of left (contralateral unloaded) knees, confirming that our mechanical loading protocol induces an OA-like histopathological phenotype. Analysis of articular cartilage lesions at one and twenty-one days post-loading show that lesions in our study are comparable to those described in a previous study at the same time point with the same loading regime [18]. Additionally, these results confirmed the spontaneous exacerbation of lesions at 3 weeks post-loading compared to lesions seen directly after loading. PTOA mice appear to exhibit pain and attempt to walk with what can be regarded as pain-related behavior. The time frame in which these lesions progress and worsen corresponds to the development of nociceptive behavior in this model, suggesting that the progressive degradation of the knee induces these profound behavioral changes. Furthermore, the mild contralateral damage we found could explain the development of the contralateral mechanical hypersensitivity seen in these animals.

Combining data from the IVIS scan for CII targeted fluorescence with the changes in mechanical properties of cartilage-bone complex for mechanically loaded mouse knees by DMA, revealed that cartilage damage is accompanied with changes in dynamic stiffness and loading energy dissipation of the complex. The results also suggest that these changes may affect the risk for damaging the subchondral bone from traumatic injury leading to OA later in life. Thus, mechanical loading increased oxidative stress, cartilage damage, subchondral bone change, and MMPs levels in the affected knee joints. The current findings will be useful as a monitoring tool for studying the progression of OA.

## Conclusions

This study helps us correlate the degree of fluorescence and the DMA value to the severity of collagen damage and subchondral bone change, respectively. These features can improve the quality of various treatment and diagnostic studies in a PTOA mouse model. Furthermore, we emphasize that the studies focused on PTOA should be comprehensive in nature and the tools provided in this study can be utilized to achieve this purpose.

In conclusion, this is the study to characterize the murine model of PTOA based on biomechanical, biochemical and behavioral parameters.

## Author contributions

HC and KAH conceived and designed the study; HC, FB, YJ and DK performed experiment and acquired the data; HC, FB, DK and DB analyzed and interpreted the data; HC, FB and DK; drafted the manuscript. HC, KAH, DB and DK proofread and critically revised it. All authors give their approval of the final submitted version.

## Funding sources

This work was supported by grants from the Arthritis Foundation (Discovery award; H. Cho) and Oxnard Foundation (Medical Research; H. Cho). This research also supported by a VA Merit Review award and VA Research Career Scientist Award (K. Hasty) from the Department of Veterans Affairs.

## Competing interest statement

The authors have no conflict of interest to declare.

## Acknowledgements

The authors want to thank Ms. Christy Patterson for her technical assistance regarding the real time rt-PCR analysis and care of animals during this study.

---

**Supplemental Figure 1.**
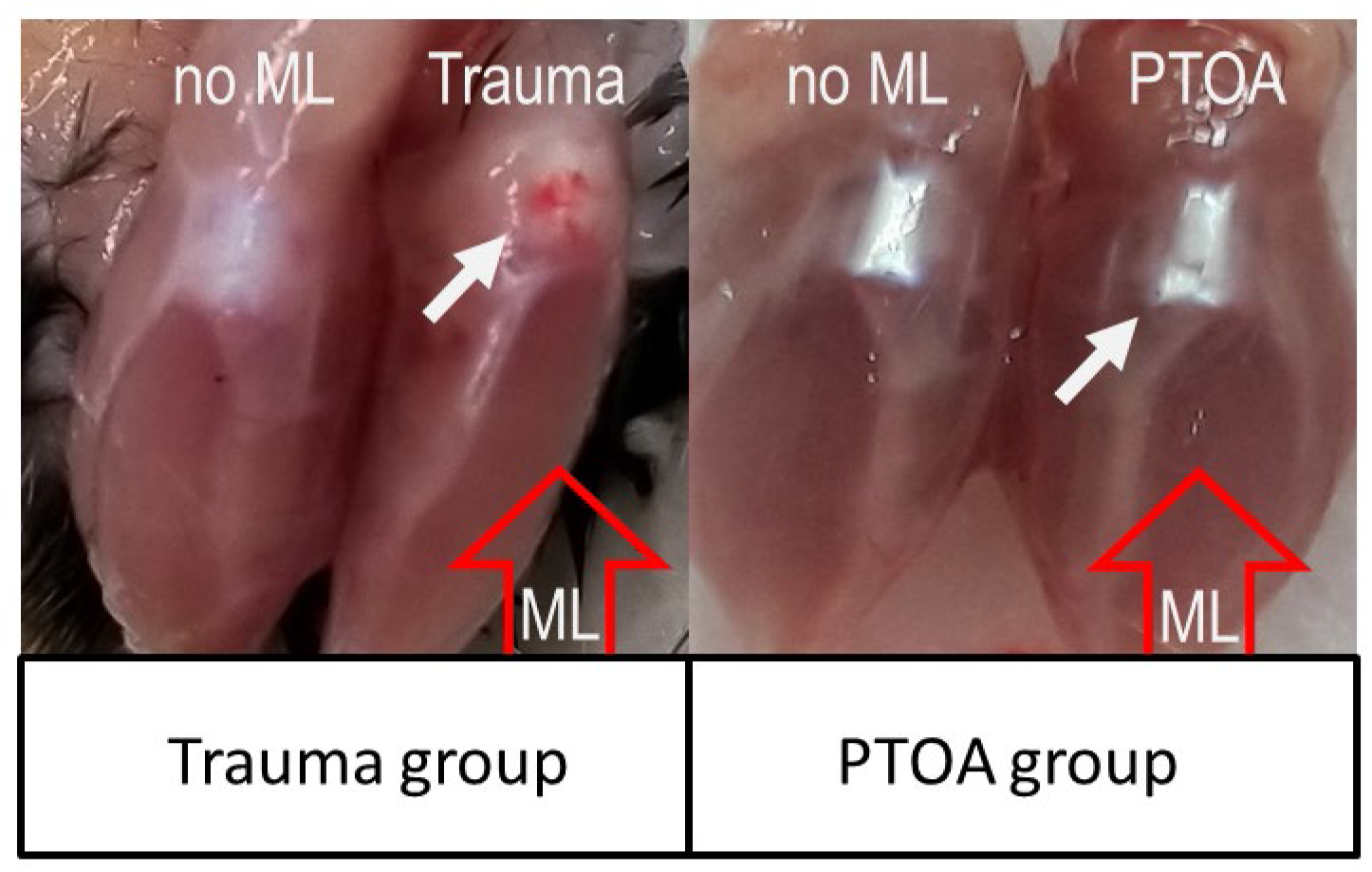
Macroscopic examination of knee joints. Blood clot formation on the knee joint in trauma group indicates severity and inflammation during joint damage. Note the disappearance of blood clot in PTOA group.

**Supplemental Figure 2.**
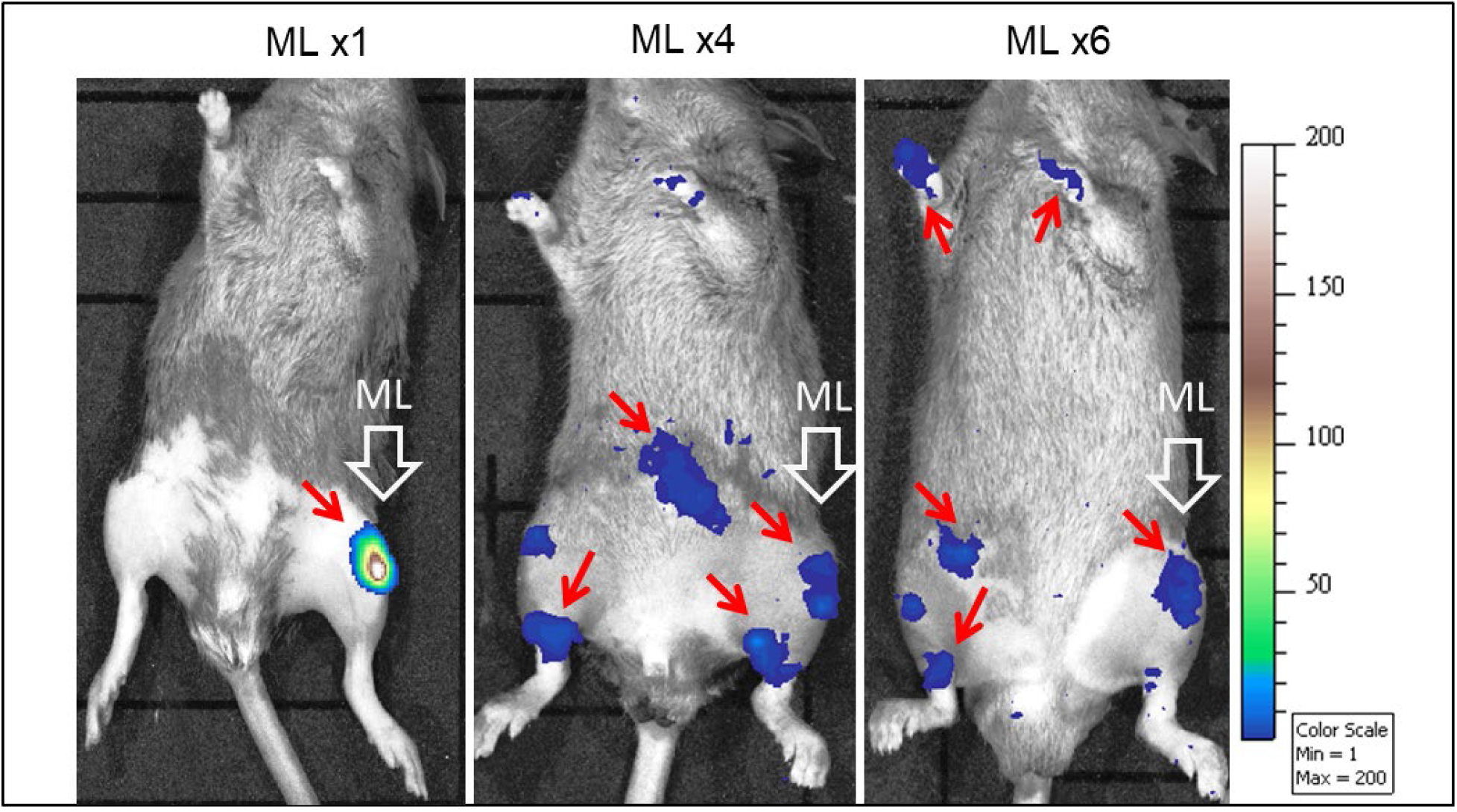
Detection of ROS-bioluminescence in knee joints of animals. L-012 is injected intraarticularly in animals after 1st, 4th and 6th mechanical loading session. Note that ROS detection is prominent only after 1st mechanical loading session. The ROS detection was not possible after successive mechanical loading sessions.

**Supplemental Figure 3.**
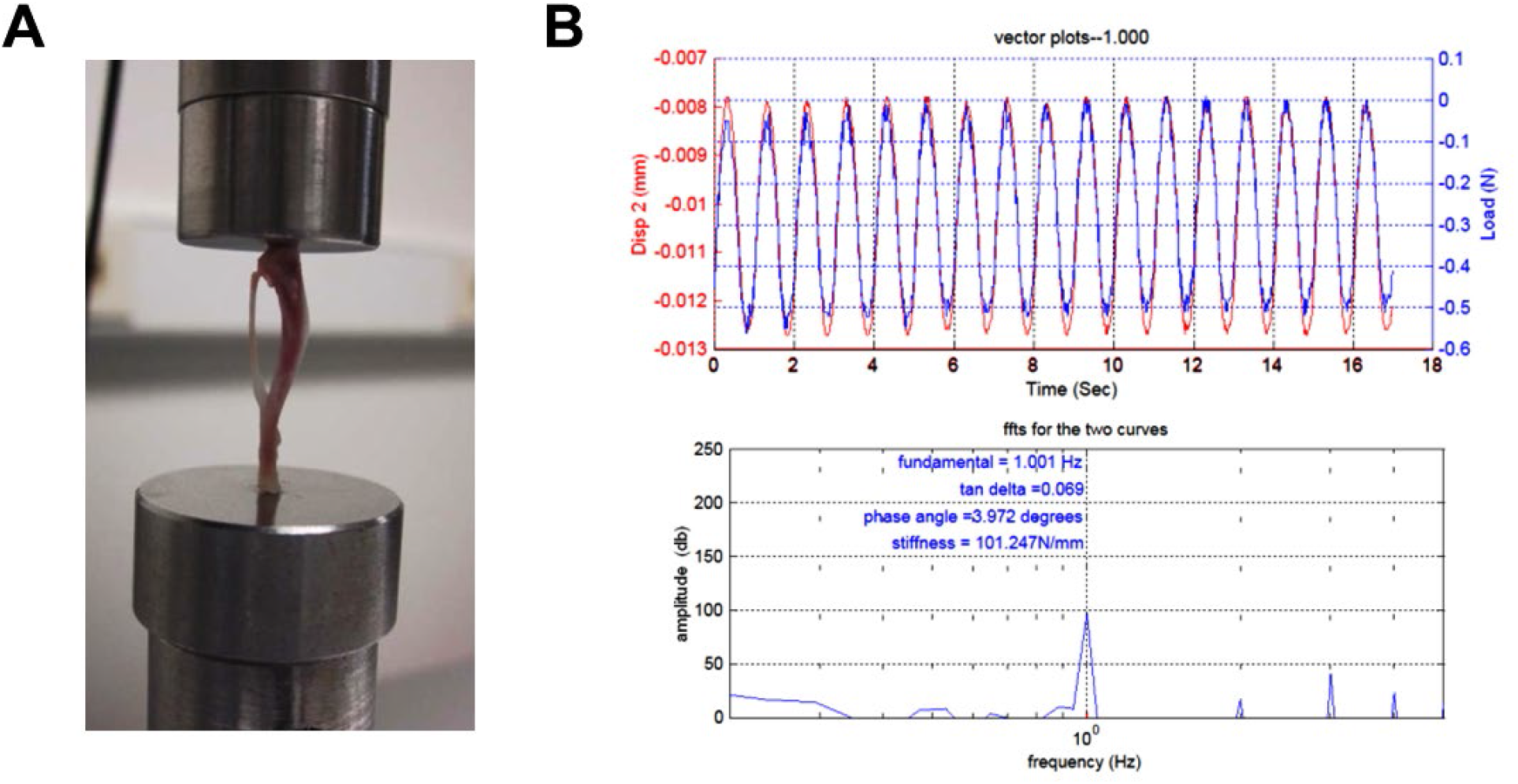
Overview of DMA (A) A mouse tibia specimen mounted on the loading machine, (B) an example of dynamic mechanical analysis (DMA) to obtain phase angle and dynamic stiffness under oscillatory loading (−0.3 ± 0.1N).

## Notes

### Competing Interest Statement

The authors have declared no competing interest.

